# Community-scale Synchronization and Temporal Partitioning of Gene Expression, Metabolism, and Lipid Biosynthesis in Oligotrophic Ocean Surface Waters

**DOI:** 10.1101/2020.05.15.098020

**Authors:** Daniel Muratore, Angie K. Boysen, Matthew J. Harke, Kevin W. Becker, John R. Casey, Sacha N. Coesel, Daniel R. Mende, Samuel T. Wilson, Frank O. Aylward, John M. Eppley, Alice Vislova, Shengyun Peng, Rogelio A. Rodriguez-Gonzalez, Stephen J. Beckett, E. Virginia Armbrust, Edward F. DeLong, David M. Karl, Angelicque E. White, Jonathan P. Zehr, Benjamin A.S. Van Mooy, Sonya T. Dyhrman, Anitra E. Ingalls, Joshua S. Weitz

## Abstract

Sunlight drives daily rhythms of photosynthesis, growth, and division of photoautotrophs throughout the surface oceans. However, the cascading impacts of oscillatory light input on diverse microbial communities and community-scale metabolism remains unclear. Here we use an unsupervised machine learning approach to show that a small number of diel archetypes can explain pervasive periodic dynamics amongst more than 65,000 distinct time series, including transcriptional activity, macromolecules, lipids, and metabolites from the North Pacific Subtropical Gyre. Overall, we find evidence for synchronous timing of carbon-cycle gene expression that underlie daily oscillations in the concentrations of particulate organic carbon. In contrast, we find evidence of asynchronous timing in gene transcription related to nitrogen metabolism and related metabolic processes consistent with temporal niche partitioning amongst microorganisms in the bacterial and eukaryotic domains.

## Introduction

Marine phytoplankton are responsible for half of global carbon fixation (*1, 2*). In tropical and subtropical marine ecosystems worldwide, most photosynthesis is attributable to cyanobacteria and picoeukaryotes, whose primary productivity is coupled to the diurnal light cycle (*3*). Beyond photosynthesis, diel patterns exist in marine microbes in transcriptional regulation of key metabolic processes such as diazotrophy, nutrient assimilation, and energy storage (*4–12*). Daily oscillations in aggregate measures of community activity (e.g., particulate organic carbon in the North Pacific Subtropical Gyre (NPSG) (*13*)) suggest that the integrated rhythms in transcription and metabolic regulation scale-up to influence biogeochemical processes such as light capture and export of matter and energy from the euphotic zone to depth. However, it remains unclear how metabolic processes occurring amongst diverse populations — including in heterotrophic bacteria that are not known to have circadian clock genes (*5*) — lead to the observed community-level dynamics.

Efforts to predict microbially-mediated ecosystem function from environmental sequence data have attempted to leverage differences in the composition of microbial metagenomes and the expression patterns within microbial metatranscriptomes as a proxy for metabolic processes. The link between genes and ecosystem function can be particularly useful when comparing different sites or seasons with significant underlying variation in diversity (*14–16*). However, there are challenges in interpreting community function from sequence data, e.g., including variable lags between transcription and translation (*17*) and unknown enzyme- and transcript-specific degradation rates (*18, 19*). In the ocean, these challenges are exacerbated by incomplete characterization of metabolic pathways and transcriptional regulatory mechanisms for many microorganisms (*20*). Furthermore, high-resolution measurements of diel patterns in biomolecule concentrations are scarce. Direct observations of biomolecule concentrations across diel cycles have revealed diurnal variation in cellular N- and C-content (*21–24*), diel rhythmicity in lipid-body formation in coral symbionts (*25*), changes in the concentrations of low molecular weight dissolved organic matter (*26*), and periodicity in intracellular metabolic products (*8, 27, 28*) in surface marine ecosystems. Studies linking diel patterns of transcription in marine microorganisms with biomolecule concentrations are typically limited to organisms in culture (*17, 29, 30*).

To explore how diel forcing of light-driven processes at the base of the marine food web affect community-level processes, we analyzed metatranscriptomes, lipidomes, and metabolomes collected every four hours over approximately three days in summer 2015 in the North Pacific Subtropical Gyre (NPSG) [http://scope.soest.hawaii.edu/data/hoelegacy/] (Figure 1A/B). This high multi-omics sampling resolution is complementary to efforts to survey for microbial diversity at the broadest ocean scales (*15, 31–33*) and to time series studies meant to infer associations within complex microbial communities at monthly/seasonal scales (*34–36*). Recent fine-scale temporal resolution studies have revealed robust diel patterns in transcriptome oscillations within complex assemblages of both photoautotrophs and heterotrophic bacteria (*5, 9, 36–38*). Here, we synthesized transcriptional data of Bacteria and Eukarya, as well as biomolecule data, and used a combination of time series analytics and machine-learning methods to examine regularity, clustering, and (a)synchronization in transcriptional activity amongst diverse taxa as well as in aggregate indicators of community metabolism.

**Fig. 1.**
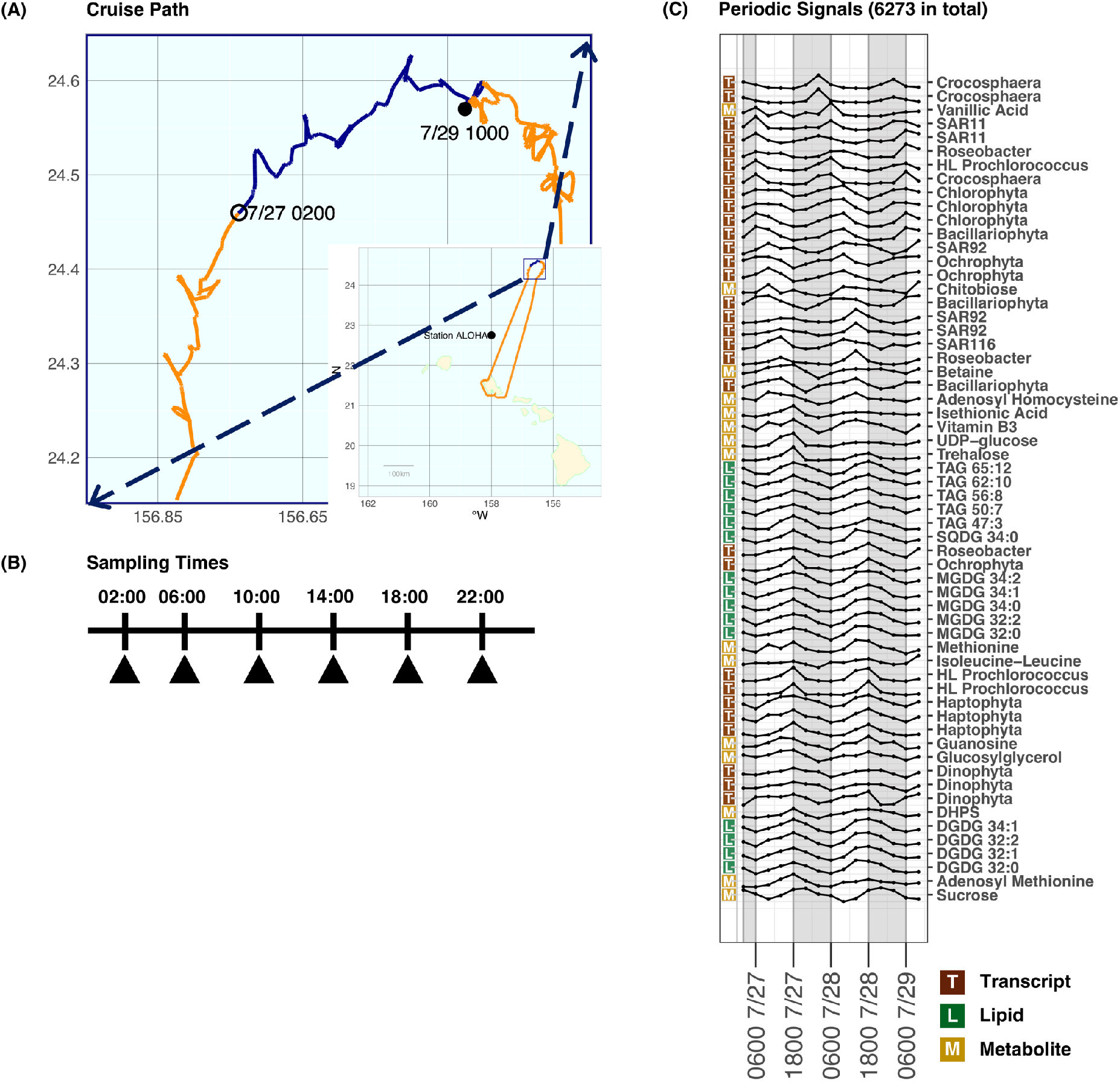
Diel patterns in diurnally resolved multiomics at Station ALOHA. (A) Map showing Lagrangian cruise track of HOE Legacy IIA cruise (orange line) with the samples used for this study taken over a 3 day period (blue line). (B) Sampling times; 4 hrs between samples.(C) Selected periodic signals determined by non-parametric analysis (see methods) ordered by peak time. Subset denotes those with lowest p-values, representing a combination of particulate metabolites (M), lipids (L), and transcripts (T) from both the >0.2 μm size fraction (predominantly prokaryotic organisms) and the >5 μm size fraction. All measurements are scaled to have mean 0 and variance 1. Grey boxes indicate nighttime hours (1800-0600).

## Pervasive Diel Periodicity in Cross-Domain Transcription, Metabolites, Lipids, and Macromolecules

We first set out to determine the extent of diel periodicity within time series of transcriptomes, metabolites, and lipids. We leveraged gene reference databases tailored to marine microorganisms (*39, 40*) to taxonomically classify transcripts from both >0.2 μm size-fraction and >5 μm size-fraction metatranscriptomes (Methods). To facilitate comparisons across taxa, a subset of transcripts with KEGG orthology annotations (*41*) were used as the basis for subsequent analyses. In parallel, we used targeted metabolomics to measure the concentrations of particulate metabolites, including free amino acids, saccharides, and vitamins. We also used lipidomic methods to measure six different classes of molecules: cell membrane-related lipids (phospholipids, betaine lipids), triacylglycerols (TAGs), chloroplast membrane-related lipids (mono- and digalactosyldiacylglycerols, sulfoquinovosyldiacylglycerols), pigments, carotenoids, and quinones (Methods). In total, we tested for diel periodicity amongst 997 unique lipids, 77 metabolites, total hydrolysable amino acids (THAA), total hydrolysable nucleobases (THNB) and 64,011 transcripts. The transcripts mapped to 5,540 KEGG orthologues from 27 major prokaryotic clades in a metatranscriptome sequenced from a >0.2 μm size fraction, and 58,471 KEGG orthologues across a selected list of 14 major eukaryotic phyla (see Methods) sequenced from a > 5 μm size fraction. We assessed periodicity using a non-parametric method (*42*), accounting for non-stationarity and multiple testing (Methods).

We identified 6,273 time series with statistically significant diel rhythms out of more than 65,000 time series examined. These significantly diel time series encompassed 501 lipids (50.2%), 50 metabolites (64.9%), 1,739 of the > 0.2 μm fraction transcripts (31.3%), and 3,983 of the > 5 μm fraction transcripts (6.8%) (see Figure 1 for represented subset of diel time series from each data set). Diel rhythms were identified in all tested taxa in both size fractions, all classes of lipids measured, and both primary and secondary metabolites, as well as in macromolecular measurements of THAA and THNB (Supplementary Files 1 & 2). In addition to metabolites and macromolecules shared across all microbial groups, we identified diel transcriptional patterns in more than 3,000 unique KEGG orthologues, spanning functions that include photosystems, photosynthetic carbon fixation, and central carbon metabolism as well as macro- and micronutrient and metal uptake (Supplementary file 1).

## Unsupervised Learning Approach to Partition and Interpret Diel Signal Patterns

The cumulative set of 6,273 diel periodic signals differed in amplitude, shape of oscillation, and peak timing. In order to reduce complexity, we implemented an unsupervised, self-organizing map (SOM) approach to determine the extent to which the data could be represented by a far smaller set of archetypes, each with its own characteristic temporal signature (Methods). SOM analysis revealed that diel signals robustly cluster into four archetypal time series with peaks at dusk (1800 hrs), night (0200 hrs), morning (0600 hrs), and afternoon (1400 hrs) (see Figures 2a and 2b; analysis of clustering resolution in Supplementary File 3). Notably, this clustering did not rely on *a priori* assumptions of sinusoidal patterns nor of preferred phase. Of the total signals, the ‘dusk’, ‘night’, ‘afternoon’, and ‘morning’ clusters comprised 36%, 22%, 18%, and 24% of the signals, respectively (Supplementary File 4). Archetypal diel clusters were heterogeneous in analyte type and taxonomic identity (i.e., each including bacterial and eukaryotic transcripts, lipids, and metabolites, Figure 2c and Supplementary Files 3).

**Fig. 2.**
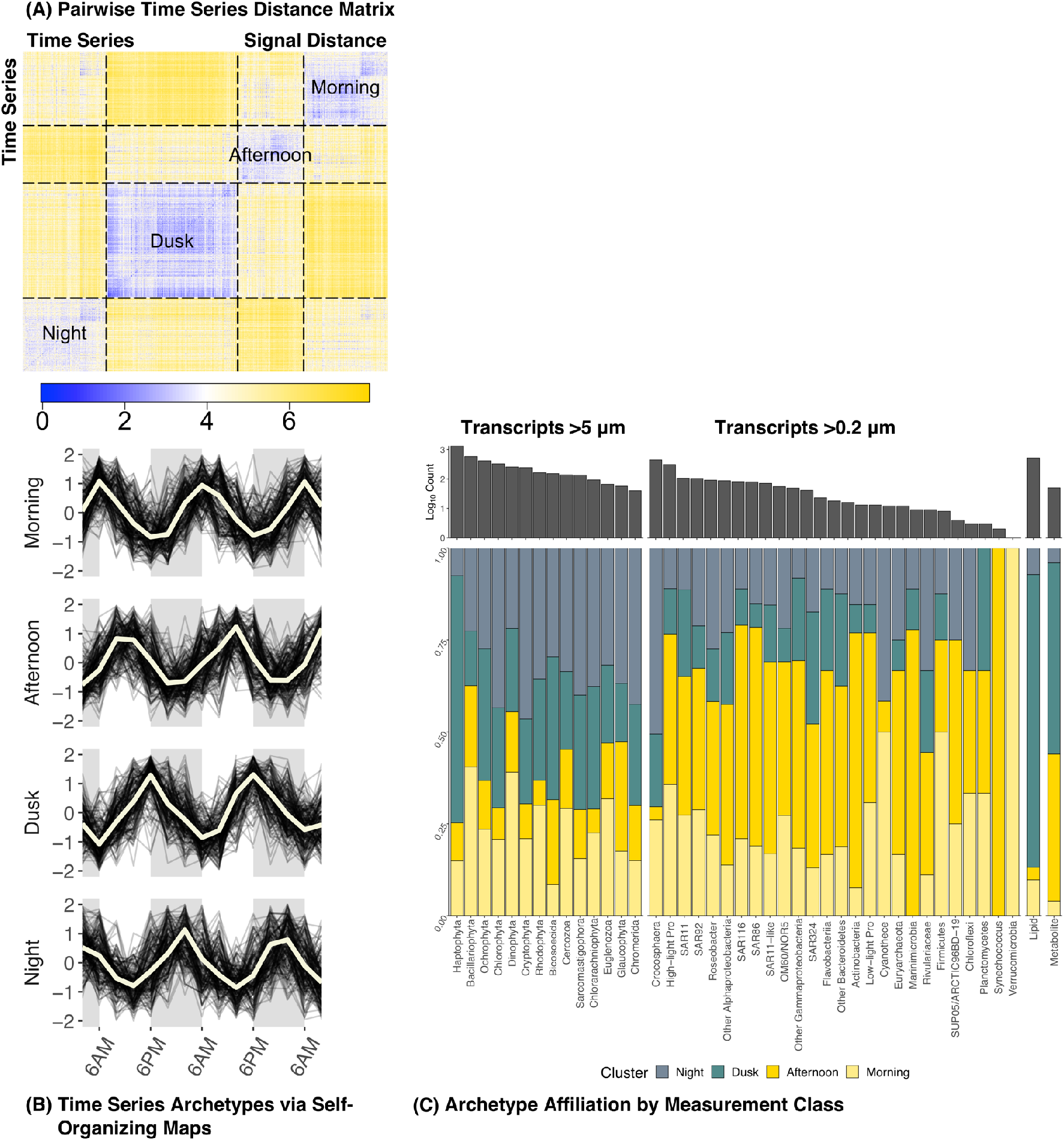
Unsupervised clustering analysis categorized diel patterns into four potential archetypes. (A) Organized pairwise distance matrix for all diel measurements after clustering based on self-organizing maps. Each pixel represents the Euclidean distance between the time series of two diel measurements, blue indicating small distance (similar time series), yellow indicating larger distance (less similar time series). Boxes are drawn around cluster boundaries. (B) Archetypal time series for four clusters (beige lines), archetypes are a combination (determined via the self-organizing map algorithm) of all time series in their cluster. A random sample of 200 time series belonging to each cluster are plotted as dark lines. (C) Distribution of diel signals across clusters for transcripts assigned to taxa from (left to right) the >5 μm transcriptome, >0.2 μm transcriptome, lipidome, and metabolome. The corresponding bar chart above indicates the quantity of signals found to be diel belonging to each group (note log scale).

We implemented a pathway enrichment analysis to summarize the distributions of transcripts for metabolic pathways among the four SOM clusters (Methods). Briefly, diel transcripts were split among three broad categories: cyanobacteria, heterotrophic bacteria, and eukaryotes. Within each category, diel transcripts were grouped by their assigned KEGG pathway. Fisher’s Exact Tests were used to identify KEGG pathways which were significantly overabundant in any of the four SOM clusters (Supplemental File 7, see Box 1 for extended results and discussion). We found significant enrichments for cyanobacterial and eukaryotic photosynthesis transcripts in the morning cluster, as found previously across oceanic ecosystems (*4, 5, 37, 38*). We also found evidence for synthesis of nucleoside and amino acid precursors, carbohydrates, and carbon fixation in the afternoon, carbohydrate catabolism at dusk, and *de novo* protein synthesis at night (see Box 1). Notably, we found indications of synchronized diel responses by heterotrophic bacteria, e.g., enrichment of the tricarboxylic acid (TCA) cycle pathway in the afternoon consistent with an increase in organic carbon catabolism given the accumulation of fixed carbon by photosynthetic organisms. We found evidence for protein synthesis occurring overnight for many eukaryotes and cyanobacterial photoautotrophs, supporting a long-standing hypothesis of staggered, diel patterns in resource acquisition, allocation, and cell division (*43*).

Altogether, we observed a synchronized cascade across diverse bacterial and eukaryotic photoautotrophs from photosynthesis in the morning and afternoon to the accumulation of organic storage molecules at dusk. The accumulated carbon provides energy for respiration and a transition to protein synthesis overnight, culminating in the synthesis of photosynthetic machinery and pigments for the upcoming dawn. We also identified a synchronized feedback by heterotrophic bacteria to the primary productivity cascade (see Figure 3). Specifically, we found that heterotroph expression of sugar uptake transporters coincides with the decline of particulate stocks of sugars and lipids. The heterotroph sugar transporters, further supported by diel oscillations in the TCA cycle pathway amongst heterotrophs (see Figure 3), suggest that cascades through the marine food web across broad domains are light controlled, as has been hypothesized previously (*37, 38*). Motivated by these results, we sought to explore the diversity within these broad taxonomic designations and to examine specific metabolic functions which may be overlooked in pathway-level analysis.

**Fig. 3.**
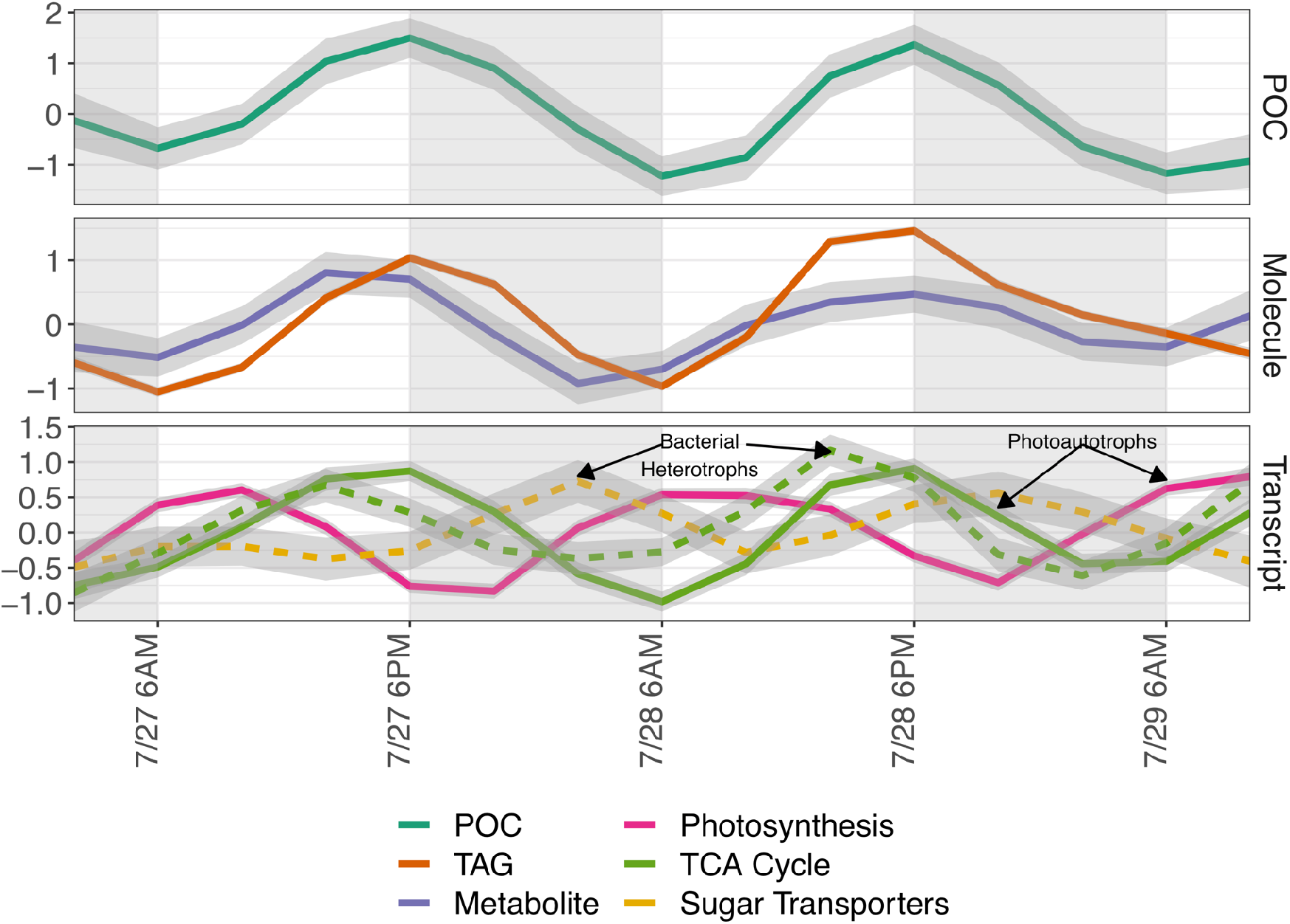
Carbon-related transcriptional and biogeochemical activity at the community scale. Averaged z-score scaled time series of optically-derived particulate organic carbon (POC) concentration (data presented in (*13*)), concentration of carbon-fixation associated lipids (and transcript levels of transcripts involved in the photosynthesis and tricarboxylic acid (TCA) cycle KEGG pathways, as well as transcripts with sugar transporter function assigned to heterotrophic bacteria. Dashed lines indicate transcripts assigned to heterotrophic bacteria. Data in these panels are smoothed using a generalized additive model with cubic spline smoothing with shrinkage penalties on all observations (95% confidence interval shown in shaded area).

## Fine-Grained Diel Transcriptional Dynamics Among Taxonomic Groups

To compare diel transcriptional patterns between taxa, we projected the aggregate time series of diel signals into a lower-dimensional space using non-metric dimensional scaling (NMDS; see Methods). The NMDS projection highlights the differences in overall transcriptional patterns between taxa and naturally projects the data into a 24-hr ‘clock-like’ space (see Figure 4). For example, Haptophytes (a phylum of algae) transcribe a higher proportion of genes with diel expression at dusk, the diazotrophic cyanobacteria *Crocosphaera* transcribe many diel genes at night, and the genes of the cyanobacteria *Prochlorococcus* with diel transcription tend to peak in the morning. Taxon-specific diel transcriptional peaks occur throughout the day and this differential peak expression shows characteristic ‘profiles’ when comparing amongst different taxa (chi-squared test of homogeneity, df=135, p<1e-5).

**Fig. 4.**
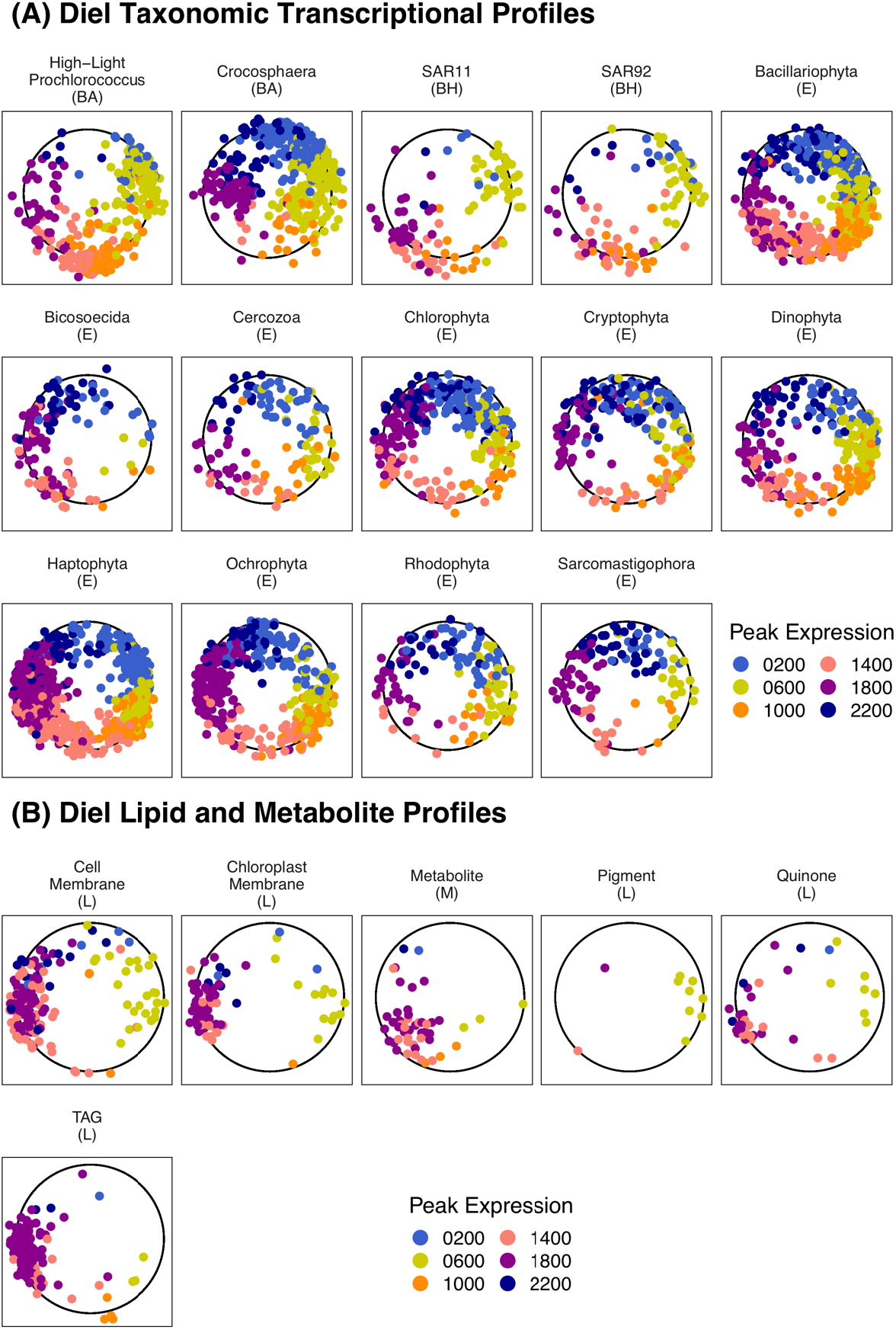
Non-metric multidimensional scaling (NMDS) projection of time series for each diel measurement in the transcriptomes, lipidomes, and metabolomes. (A) Each point represents one gene with diel transcription. Diel transcriptional peaks are distributed around an emergent 24-hour ‘clock’ and amongst community members and metabolic functions. Points are colored by calculated peak rank measurement time (see Methods). Parentheses indicate taxonomic affiliation: (E) – Eukaryote; (BH) – Bacteria Heterotroph; (BA) – Bacteria Autotroph. Only taxa with at least 100 diel transcripts are shown. (B) Projections of diel metabolites (M) and lipids (L). Each point represents one metabolite or lipid. Lipids are separated by functional categories.

Next, we analyzed the oscillatory patterns of diel genes to explore the potential functional basis underpinning the differences between the taxonomic profiles. KEGG orthologues were classified into three categories: (1) ‘taxonomically narrow’ diel transcripts, corresponding to KEGG orthologues that only show diel transcriptional patterns in 3 or fewer taxa, (2) ‘synchronous’ diel transcripts, corresponding to KEGG orthologues that show diel expression in more than 3 taxa and have a uniform diel transcriptional pattern (i.e., the expression peaks at similar times); and (3) ‘asynchronous’ diel transcripts, corresponding to KEGG orthologues that are expressed in more than three taxa but have discordant diel transcriptional patterns (i.e., the expression peaks at different times). Of the 3,193 KEGG orthologues which showed a diel transcriptional pattern in any taxon, the vast majority (2,898) was taxonomically narrow, i.e., a diel signature was only observed in three or fewer taxa (Supplemental File 7). Hence, characteristic taxonomic profiles (Figure 4) are driven primarily by transcripts that have diel expression in a small set of taxa in our dataset. This may point to taxon-specific differences in regulation and functional capacity (Supplemental File 9). The remaining 295 widely shared KEGG orthologues were classified as synchronous or asynchronous based on the average difference in peak time across all associated taxa for that KEGG orthologue relative to a null expectation (Methods).

We identify 80 of 295 orthologues as having ‘synchronous’ diel transcription (BH adjusted p-value<0.1) (Figure 5, Supplemental File 7). The synchronous orthologues with the most associated diel signals include photosystem II (PSII) components and cytochrome C oxidases. Cytochrome C oxidases are found in the photosynthetic e chain as well as in oxidative phosphorylation in both photosynthetic and non-photosynthetic organisms (e.g., heterotrophs). The widespread diel synchronicity of these orthologues is consistent with the hypothesis that biophysical and/or regulatory constraints transcend taxa in dictating peak time for functions related to photosynthesis and oxidative phosphorylation (Figure 3); as seen in other community-level studies (*37*). Hence, we interpret this evidence to imply that the emergent primary productivity cascade is driven, in part, by a few highly conserved genes with widespread synchronous diel expression.

**Fig. 5.**
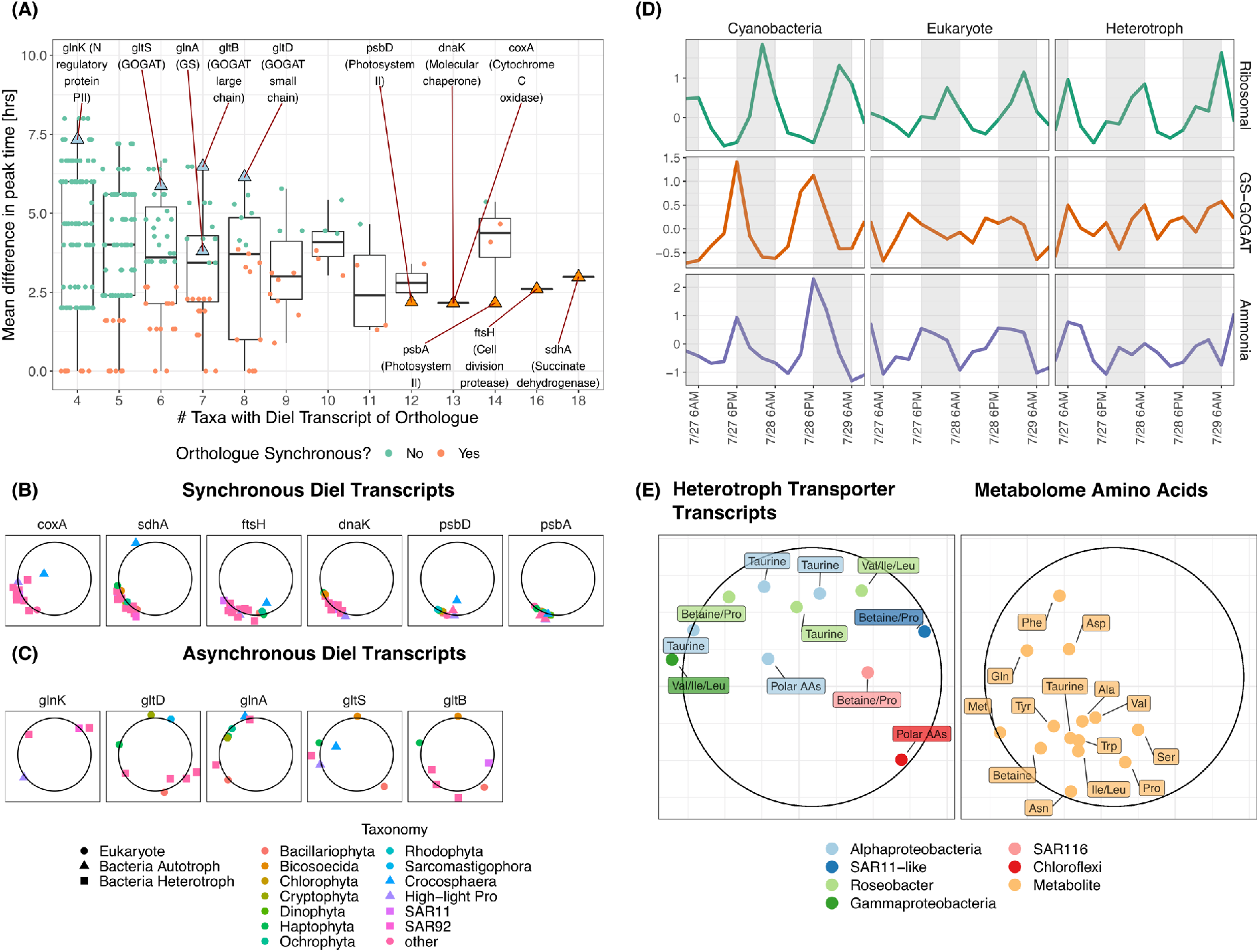
Niche partitioning and nitrogen metabolism at Station ALOHA. **(A)** For each KEGG orthologue with diel expression in at least 4 different taxa, all pairwise differences in peak rank time were tabulated and averaged (see Methods). Low average difference in peak time indicates that most taxa with diel transcription of that orthologue peaked at the same time of day. Orthologues with smaller average differences in peak times than would be expected from the population of all diel transcripts with an BH-adjusted p-value <0.1 are indicated in orange. Orthologues related to the GS-GOGAT system are labeled and indicated as green triangles. Orthologues with low average peak time difference expressed widely, related to primary production and central carbon metabolism are labeled as orange triangles. (B) NMDS projection for the subset transcripts for synchronous genes indicated in (A). Taxonomic designations are indicated by color and shape. (C) NMDS projection of the subset of transcripts for the asynchronous GS-GOGAT related genes indicated in (A). (D) Transcription dynamics of ammonia transporter (amt), GS-GOGAT, and ribosomal subunit genes over eukaryotes, cyanobacteria, and heterotrophic bacteria. Lines indicate average z-score transcription levels across all taxa with diel expression of a gene with the labeled function. Shaded boxes indicate nighttime hours. (E) NMDS projection showing peaks in expression for heterotrophic bacteria amino acid uptake transporters (left) and diel amino acids from the metabolome (right). Points are labeled with which amino acid they are, or which amino acid the transporter takes up. Color indicates taxonomic affiliation.

## Temporal Niche Partitioning and Nitrogen Metabolism

In evaluating transcripts that are ‘widespread’ (i.e., found in more than 3 taxa), we identified 25 different KEGG pathways not related to primary productivity, some of which included asynchronous diel expression patterns. An initial curation of ‘asynchronous’ orthologues with large average peak time differences included transcripts annotated as *gltB* (glutamate synthase NADPH large chain subunit) and other components of the GS-GOGAT system (Figure 5, Supplemental File 7). Glutamate synthase (part of the GS-GOGAT system) is the first intracellular step in ammonia assimilation (*44*). The expression of *gtlB* spanned both heterotrophs and autotrophs and all four SOM clusters (Supplemental File 3). Similarly, transcripts associated with other GS-GOGAT subunits, including *gltD*, *gltS*, *glnA*, and the nitrogen-regulatory PII response protein *glnK* exhibited asynchrony (Figure 5). GS-GOGAT components associated with *Prochlorococcus* peaked in the afternoon and dusk, as previously reported (*11*). Furthermore, transcripts annotated as the ammonium transmembrane transporter amt had diel expression across 10 different taxa and showed strong asynchrony. We found photoautotrophs had peak gene expression at dusk (concurrent with GS-GOGAT component transcripts), while heterotrophs had peak expression in the morning (again, concurrent with GS-GOGAT component transcripts). As such, it appears the expression of key steps required for nitrogen uptake and assimilation are periodic and asynchronous, i.e., they repeatedly peak at different times in the day across taxonomic groups. Asynchronous timing of the expression of nitrogen assimilation-related transcripts suggests microbes differ in their strategies of utilizing a key nutrient, albeit via a process which is not known to be directly linked to light. Nitrogen assimilation is of particular interest in the NPSG due to the exceedingly low background concentration of inorganic nitrogen species (nitrate, nitrite, ammonia), resulting in persistent and chronic nitrogen limitation, as well as a strong reliance on remineralization for the nitrogen supply (*45*). Dispersed nitrogen assimilation-related transcript peak times may also indicate a potential mechanism for the emergence of nitrogen stress mitigation at the community scale.

While inorganic forms of nitrogen are extremely dilute in the epipelagic NPSG, relatively more abundant organic nitrogen compounds are another important nitrogen source, especially for heterotrophs (*46–48*). We find 14 intracellular amino acids with diel periodicity (Figure 5), including the amino-sulfonic acid taurine, which has been hypothesized to be an important substrate for marine heterotrophs (*49*). We investigated if any of these diel amino acids may be of importance to heterotrophic populations by searching for corresponding uptake transporter genes with diel expression. We found that of the 71 total diel transporter genes assigned to heterotrophic bacteria, 11 take up amino acids, including leucine/isoleucine, valine, betaine, proline, and taurine (Figure 5). These molecules are all identified as diel, as is the THAA pool, indicating that they are synthesized and accumulate in cellular macromolecules on a diel cycle. In contrast, *Prochlorococcus* shows diel expression of the urea uptake transporter *urtA* in the night cluster, coincident with expression of ureases in picoeukaryotes (Supplemental Data 3). Notably, this timing follows after the peak transcription of *amt* at dusk, indicating sequential transport of ammonia and then urea overnight for these photoautotrophs. Urea is known to be among important sources of organic nitrogen for photoautotrophs in the NPSG (*50, 51*). Nighttime expression of urea uptake and catabolism genes may reflect an increase in supply due to exudation from active phagotrophic protists (*52*) or nocturnally feeding macroscopic zooplankton (*53*). In addition to evidence that uptake and assimilation of inorganic nitrogen species may be temporally partitioned among the microbial community, we also find potential niche-specific preferential uptake of diverse organic nitrogen species throughout the day.

In addition to macronutrients, our data indicate diel regulation of uptake and synthesis of micronutrients (particularly cobalamins and iron) across photoautotrophs and heterotrophic bacteria. This suggests temporal partitioning extends to diverse metabolic processes across microbial communities in the NPSG (see extended treatment in the Supplemental Information and related analysis of oscillations in the carotenoid biosynthesis pathways (*54*). Given the observation of periodicity in the uptake of organic nitrogen species, we suggest that a comprehensive accounting of community metabolism will also need to account for mortality/loss process (e.g., as caused by grazing and viral lysis) which itself exhibits periodic regularity (*52*).

## Conclusions

Our synthesis of *in situ* multi-omics revealed prevalent and previously unrecognized synchronicity in pathways and signals at Station ALOHA in the NPSG including both functions directly mediated by light (e.g., photosynthesis) and other metabolic processes which may be indirectly affected by other ecological and environmental drivers (e.g., organic substrate uptake, vitamin synthesis and exchange). Machine learning methods revealed both coherence in carbon-associated metabolism and asynchronous timing of nitrogen uptake and assimilation across the community. We hypothesize that asynchronous timing is indicative of diel niche partitioning, such that taxa may minimize competition for scarce resources such as macro- and micro-nutrients by restricting their demand for these resources to narrow time intervals via mechanisms analogous to the temporal storage effect (*55–57*). These ‘temporal niches’ may offer considerable fitness advantages to competitors for persistently scarce, periodically supplied resources. In summary, our joint analysis of both bacterial and eukaryotic transcription supplemented by direct analysis of biomolecular concentrations (of metabolites, protein, nucleic acids, and lipids) reveals how microorganisms respond directly to light input, to each other, and to their environment via the use, assimilation, and regeneration of limiting nutrients in the surface ocean. The emergence of community-scale synchronization and temporal niches reinforces the need to integrate mechanistic studies of diel cycles within complex microbial communities with global-scale models to understand the maintenance, diversity, and resilience of carbon and nitrogen cycles in future ocean scenarios.

## Acknowledgments

The authors thank Tara Clemente and Elisha Wood-Carlson for facilitating data collection. Additional thanks to the captains and crew of the R/V Kilo Moana and research staff at SOEST and Sheean Haley and Kyle Frischkorn for sample collection for eukaryotic transcriptomes.

## Author contributions

DM and JSW led writing of the manuscript. AKB, MJH, KWB, DRM, and SJB contributed to writing on the manuscript. DM, AKB, MJH, KWB, JRC, SNC, FA, AV, DRM, STW, SJB, EVA, EFD, DMK, AEW, JPZ, AEI, BASVM, STD, and JSW edited the manuscript. AKB, FA, AV, AEW, and STD contributed to data collection. AKB, MJH, KWB, FA, JE, DRM, AEI, BASVM, and STD contributed to sample processing/data preparation. STW served as chief scientist for research expedition. DM, AKB, MJH, KWB, JRC, SNC, DRM, STD, and JSW developed the data analysis methods. DM, AKB, and SNC wrote code. DM, SNC, SJB, SP, AEI, and JSW contributed to analysis design. DM, AKB, MJH, JRC, BVM, AEI, STD, and JSW analyzed data. STW, BVM, AEI, STD, and JSW designed the research, with contributions from all authors.

## Data and materials availability

Sequence data for the >0.2 μm metatranscriptome are deposited in the Sequence Read Archive through the National Center for Biotechnology Information under BioProject ID PRJNA358725. The Station ALOHA gene catalogue data are available under Bioproject no. PRJNA352737, and iMicrobe (http://datacommons.cyverse.org/browse/iplant/home/shared/imicrobe/projects/263/ALOHAgenecat_v1_nonredundant.annot). Sequence data for the >5 μm metatranscriptomes are available in the Sequence Read Archive through the National Center for Biotechnology Information under accession no. SRP136571, BioProject no. PRJNA437978. Raw files for metabolomics data are available in Metabolomics Workbench (116) under project ID PR000797. Processed data will be available soon in Metabolomics Workbench under Project ID PR000926. Lipidomics mass spectral raw data are available from authors upon request. All code and feature/abundance tables used to complete this analysis are available at https://github.com/WeitzGroup/community_scale_metabolism_NPSG archived under https://zenodo.org/badge/latestdoi/262179139.

## Supplementary Materials

**Online Box 1.** Community-Scale Synchronization of Phototrophy and Organic Carbon Catabolism.

Morning (0600): Analysis of the morning cluster revealed significant enrichment of transcripts for both cyanobacteria and eukaryotes involved in the photosynthesis pathway (Fisher’s exact test, BH-adjusted p<0.1). Eukaryotes had additional significant enrichment for transcripts in the carotenoid synthesis, carbon fixation, and porphyrin/chlorophyll metabolism pathways. The chemical data in the morning cluster corroborate transcriptional enrichments, including 9 out of 11 diel pigments from the lipidome (chlorophylls and carotenoids such as lutein and zeaxanthin) (*54, 107*) and the metabolite pantothenic acid (vitamin B_5_), which is a precursor to coenzyme A (*108*) and critical to fatty acid synthesis.

Afternoon (1400): We see evidence consistent with afternoon synthesis of amino acids, nucleosides, and carbohydates, carbon fixation, and energy metabolism-related transcriptional processes. Similar to the morning cluster, photosynthesis transcripts are enriched for both eukaryotes and cyanobacteria, and the porphyrin/chlorophyll metabolism pathway is enriched in eukaryotes. This cluster includes diadinoxanthin, a photoprotective carotenoid utilized by both dinoflagellates and diatoms (*54, 107*). Metabolites in the afternoon cluster include serine and alanine, precursors in the formation of betaine lipids produced by phytoplankton, as well as precursors involved in the synthesis of glyceroglycolipids associated with photosynthetic carbon fixation, such as chitobiose and UDP-glucosamine. This cluster also shows significant enrichments of the TCA cycle and oxidative phosphorylation pathways in both eukaryotes and heterotrophic bacteria, indicating that these groups ramp up organic carbon catabolism in response to the production of fixed carbon. Additionally, heterotrophic bacteria have enrichments for transcripts in the RNA degradation and cell cycle pathways (primarily proteases and molecular chaperones), potentially indicating cell replication. Our findings that bacteria respond during the day to light-driven forcing is consistent with prior evidence of increased bacterial production during the daytime at Station ALOHA (*109*).

Dusk (1800): The dusk cluster shows transcriptional and chemical evidence of the cumulative effects of carbon fixation and transition to catabolism and replication with sunset. Over 90% of diel triacylglycerol lipids (TAGs), storage lipids used by eukaryotic phytoplankton (*8*), and over half of the diel metabolites fall into this cluster. These metabolites include several osmolytes known to be produced by prokaryotic and eukaryotic phytoplankton (*27, 110*), such as the saccharide trehalose and organosulfur molecules dimethylsulfoniopropionate (DMSP) and 2,3-dihydroxypropane-1-sulfonate (DHPS) (*28*). The dusk enrichment of tricarboxylic acid (TCA) cycle transcripts in both cyanobacterial and eukaryotic groups suggests the use of this accumulated organic carbon as an energy source. We also find an enrichment in eukaryotic transcripts associated with fatty acid oxidation to acetyl-CoA, particularly in haptophytes and ochrophytes. This corresponds with the peak in concentrations of TAGs and the fatty acids arachidonic acid and eicosapentaenoic acid (EPA), the latter of which has been shown to be a major constituent of the fatty acid profiles of the betaine lipids of eukaryotic phytoplankton in subtropical gyres (*111*). Over 75% of the diel betaine lipids (55/73) and phospholipids (87/116) peak at dusk, following the peak of many of their precursor metabolites in the afternoon. These major classes of polar glycerolipids are important constituents of the cell membranes of eukaryotic phytoplankton (*112*), and biological membranes of all marine organisms, respectively. The dusk cluster encapsulates the peak in concentration of carbohydrates and other products of primary production in the particulate fraction, as well as transcripts suggesting the breakdown of these products across the microbial community for energy.

Night (0200): The night cluster contains transcriptional patterns that indicate a shift in community-wide metabolic processes away from the synthesis of lipids and secondary metabolites e.g., including only 7.8% of diel lipids and 4% of diel metabolites). Both eukaryotes and cyanobacteria show significant enrichment for ribosomal subunit transcripts, suggesting an increase in protein synthesis. This shift is corroborated by the nighttime peak in total hydrolysable amino acids and total hydrolysable nitrogenous bases. In the night cluster we also find transcripts involved in nitrogen fixation in the cyanobacterium *Crocosphaera* (simultaneous data reported in 6), including nitrogenase and accessory proteins such as hydrogenase, ferrous iron transporters, and superoxide dismutase. This cluster also contains *Crocosphaera* transcripts for the biosynthesis of nitrogen-rich molecules such as pseudocobalamin (Supplemental Text) and chlorophyll, which is reflected in the presence of pigments in the morning cluster. Diel expression for the biosynthesis of nitrogen-rich secondary metabolites also extends to nonribosomal peptide synthetases in eukaryotes. Similarly, heterotrophic bacterial transcripts related to the nitrogen-demanding porphyrin/chlorophyll metabolism and photosynthesis antenna protein pathways were significantly enriched at night. Further inspection revealed overnight diel enrichment of the complete biosynthetic pathway from protoporphyrin IX to bacteriochlorophyllide a in the Roseobacter group of Alphaproteobacteria and the OM60/NOR5 group of Gammaproteobacteria, both of which synthesize this pigment (*113, 114*). In summary, we find evidence of nighttime production of proteins associated with cellular division, nitrogen rich compound production, and the preparation of photoheterotrophic machinery before sunrise.

## Materials and Methods

### Fieldwork design and sampling

Fieldwork was conducted during 25 July to 5 August, 2015 in the oligotrophic North Pacific Subtropical Gyre. To maximize the signal to noise ratio in an open ocean environment, a Lagrangian sampling strategy was implemented whereby World Ocean Circulation Experiment Surface Velocity Profile (WOCE SVP) drifters from Pacific Gyre, Inc. were deployed within the center of a mesoscale anticyclonic eddy. The mesoscale eddy fields were identified using Archiving, Validation, and Interpretation of Satellite Oceanographic data (AVISO) and when the field sampling occurred, the target anticyclonic eddy was located north of the Hawaiian Islands at 24.4 N and 156.5 W, with a diameter of ~100 km. Over the 12-day sampling period, the shipboard measurements were conducted alongside the drifters as they performed an almost complete circular pattern with a diameter of ~44 km (Figure 1a). Water-column seawater sampling for diel measurements took place every 4 h for a period of 4 days (26-30 July) and 3 days (31 July-3 August) at a depth of 15 m corresponding to the depth of the drogue. The water-column sampling was achieved using a 24 × 12 L Niskin bottle rosette attached to a conductivity-temperature-depth (CTD) package (SBE 911Plus, SeaBird) with additional fluorescence, oxygen, and transmissometer sensors. The sampling and analytical protocols for vertical profiles of nutrients, particulates, and flow-cytometry enumerated phytoplankton populations and heterotrophic bacteria were identical to those employed by the Hawaii Ocean Time-series program (http://hahana.soest.hawaii.edu/index.html). A more detailed explanation of the sampling strategy and resulting datasets can be found in Wilson et al. (*1*).

### Metabolite sample collection, extraction, and analysis

Metabolite data was collected as described previously (*2, 3*). Briefly, 3.5 L of seawater was filtered onto a 47 mm, 0.2 Omnipore filter using a peristaltic pump and flash frozen in liquid nitrogen. Samples were collected in triplicate at every time point. Filters were stored in a −80 °C freezer until the time of metabolite extraction. Metabolites were extracted as previously reported (*3*) with a modified Bligh and Dyer extraction (*4*) using 1:1 methanol:water (aqueous phase) and dichloromethane (organic phase) to extract aqueous and organic metabolites. Select isotope labeled internal standards were added before or after extraction to aid in normalization (*5*). Metabolites were measured with a Waters Xevo TQ-S triple quadrupole and a Thermo Scientific Q-Exactive Orbitrap HF with both reversed-phase and hydrophilic interaction liquid chromatography (HILIC). Metabolite peaks were integrated with Skyline for small molecules (*6*), followed by quality control and normalization. Details of the data acquisition and processing have been previously reported (*3, 5*). Blank filters were extracted alongside the samples. Metabolites that did not pass quality control in more than 10% of samples were discarded, further discussed in Boysen et al 2020 (*3*). For metabolites that passed the quality control in 90% of samples but not all samples, the remaining samples were filled in with values to reflect the limit of detection for that metabolite.

### Macromolecular Measurements

Macromolecules were hydrolyzed as in Fountoulakis and Lahm (*7*) with some modifications as follows: Samples were heated at 120 °C for 20 hours instead of 110 °C for 20-24 hours since initial recovery tests with bovine serum albumin (BSA) resulted in better recovery of the amino acids at 120 °C as compared to 110 °C or a shorter hydrolysis with BSA at 150 °C. Punches of 142 mm 0.2 μm Durapore filters were transferred into acid (10% hydrochloric) and solvent (water, methanol, dichloromethane) cleaned 40 mL teflon centrifuge tubes. Enough 6N hydrochloric acid was added to cover the filter along with spikes of isotope labeled amino acid and nucleobase standards. Each sample was purged under nitrogen gas for 30 seconds before immediately being sealed with a solvent rinsed cap. The samples were heated at 120 °C for 20 hours. The acid was then transferred to a clean, combusted glass vial. The original teflon vial and filter were rinsed with approximately 500 μL of optima grade water and transferred to the new glass vial. A rinsing step was repeated with an equal volume of optima grade methanol. The acid mixture was concentrated to dryness under nitrogen gas and on a heat block set to medium heat. Once dried, approximately 500 μL of water was used to rinse each vial and samples were returned to dry completely under the nitrogen gas. Dried samples were re-dissolved in 1 mL of optima grade water and syringe filtered into LCMS vials.

LC-MS analysis of nucleobases and amino acids used a SeQuant ZIC-pHILIC column (5 μm particle size, 2.1 mm × 150 mm, from Millipore) with 10 mM ammonium carbonate in 85:15 water to acetonitrile (Solvent A) and 10mM ammonium carbonate in 85:15 acetonitrile to water (Solvent B) at a flow rate of 0.15 mL/min. The column was held at 100% B for 2 minutes, ramped to 64% A over 18 minutes, ramped up to 100% A over 1 minute, held at 100% A for 7 minutes, and equilibrated at 100% B for 22 minutes (total time is 50 minutes). The column was maintained at 30 °C. Compounds were detected on a Thermo Scientific Q-Exactive Orbitrap HF with a full scan method employing positive and negative switching, a scan range of 60 to 900 m/z, and a resolution of 60,000. The capillary temperature was 320°C, the H-ESI spray voltage was 3.5 kV, and the auxiliary gas heater temperature was 90°C. The S-lens RF level was 65. Sheath gas, auxiliary gas, and sweep gas flow rates were maintained at 16, 3, and 1, respectively.

### Lipidome

Detailed methods for lipidomics are described in Becker et al (*8*). In brief: Lipids were extracted from triplicate samples using a modified Bligh and Dyer protocol (*9*). The total lipid extract was analyzed on an Agilent 1200 high performance liquid chromatography (HPLC) system coupled to a ThermoFisher Exactive Plus Orbitrap high resolution mass spectrometer (HRMS; ThermoFisher, Waltham, MA, USA) equipped with an electrospray ion source. Analyte separation was achieved using reversed phase HPLC on a C8 Xbridge column (particle size 5 μm, length 150 mm, width 2.1mm; Waters Corp., Milford, MA, USA). HPLC and MS conditions are described in (*10*) (modified after (*11*)). For the identification and quantification of lipids, we used LOBSTAHS, an open-source lipidomics software workflow based on adduct ion abundances and several other orthogonal criteria (*10*). Lipids identified using the LOBSTAHS software were quantified from MS data after pre-processing with XCMS (*12*) and CAMERA (*13*) and corrected for response factors of commercially available standards as described in (*8*) and Biller et al. (under review). As a means of validating the accuracy and reliability of LOBSTAHS identification and quantification, quality control (QC) samples of known composition were interspersed with the environmental samples.

### >0.2 μm metatranscriptome

The >0.2 μm transcriptomes were collected, sequenced, quality controlled, quantified, and normalized, as described previously (*1, 14*). Briefly, 2 L of seawater was filtered onto 25 mm, μm Supor PES Membrane Disc filters (Pall) using a peristaltic pump. The filtration time was between 15 and 20 min., and filters were placed immediately in RNALater (ThermoFisher Scientific, Waltham, MA) and preserved at −80 °C until processing. RNA extractions were performed by removing RNALater (ThermoFisher Scientific, Waltham, MA) followed by the addition of 300 μl of Ambion denaturing solution directly to the filter, and then vortexing for 1 min. Next, 750 μl of nuclease free water was added, and the samples were robotically purified and DNase treated using the Chemagen MSM I instrument with the tissue RNA CMG-1212A kit (Perkin Elmer, Waltham, MA). RNA quality was assessed using Fragment Analyzer high sensitivity reagents (Advanced Analytical Technologies, Inc.), and quantified using Ribogreen (Invitrogen, Waltham MA).

Internal standard RNA mixtures used for quantitative transcriptomics were prepared as previously described (*1, 14*). Internal standards were added at known concentrations to samples prior to sequencing by a Nextseq500. Generated reads were trimmed of adapter sequences with Trimmomatic v 0.27 (*15*), end-joined with PandaSeq v2.4 (*16*), and filtered for quality using Sickle v1.33 (*17*). Reads containing ribosomal RNA sequences were removed in silico using sortmerna v2.1 (*18*). Spiked-in RNA internal standard sequences were identified using lastal v756 (*19*), quantified, and then removed. The remaining transcript reads were mapped to the merged HOE Legacy II-ALOHA metagenomic gene catalogue using lastal, as previously described (*1, 14*). For each time-point, the average normalization coefficient (derived from five different internal standards) was multiplied by the reads mapped to each transcript, to estimate transcripts per liter for each gene in the sample. This normalized transcript count table, and HOE Legacy II-ALOHA metagenomic gene catalogue annotations (*1, 20*), were used in subsequent bioinformatic analyses of the >0.2 μm sample transcripts. The >0.2 μm metatranscriptome data are deposited in the Sequence Read Archive through the National Center for Biotechnology Information under BioProject ID PRJNA358725. The Station ALOHA gene catalogue data are available under under Bioproject no. PRJNA352737, and iMicrobe (http://datacommons.cyverse.org/browse/iplant/home/shared/imicrobe/projects/263/ALOHAgenecat_v1_nonredundant.annot).

### >5 μm metatranscriptome

The >5μm metatranscriptomes were collected, sequenced and quality controlled as previously described (*21*). Briefly, for each time point, 20 L of water was filtered onto two 5 μm 47 mm polycarbonate filters by way of peristaltic pump, passing ~10 L through each filter. Filtering time did not exceed 40 minutes, upon which filters were placed into liquid nitrogen until processing. RNA extractions were performed using a Qiagen RNeasy Mini Kit (Qiagen, Hilden, Germany), modifying the lysis step with the addition of Biospec 0.5 zirconia/silica beads. For each filter set (n=2, representing 20 L of sample volume), lysis buffer and beads were added, vortexed for 1 minute, placed on ice for 30 s, and vortexed again for 1 min. Lysate was removed with a pipette and pooled into a single 5 ml microcentrifuge tube. The rest of the Qiagen RNeasy Mini Kit protocol was then followed according to the manufacturer’s instructions, adjusting volumes accordingly and incorporating the on-column DNase digestion step, using a Qiagen RNase-free DNase kit. Resulting total RNA was eluted with RNase-free water and then purified and concentrated with a RNeasy MinElute kit according to the manufacturer’s instructions. Quantity and quality of extracted total RNA were assessed on an Agilent 2100 Bioanalyzer (Agilent, Santa Clara, CA). Illumina Truseq libraries were prepared at the JP Sulzberger Columbia Genome Center following center protocols and sequenced on an Illumina HiSeq 2000 to produce 90 million 100-bp, paired-end poly-A selected reads. Raw sequence quality was visualized, and then cleaned and trimmed as previously described (*21*).

Mapping of the >5 μm metatranscriptome was conducted using the Burrows-Wheeler Aligner (BWA-MEM, parameters -k 10 -aM; (*22*)) against a reference database constructed from MMETSP transcriptomes after Alexander et al. (*23*). Resulting alignments were counted using the HTSeq 0.6.1 package (options -a 0, --m intersection-strict, -s no; (*24*)). Read counts were then filtered for contigs with average read counts ≥ 10 across the time series and then DESeq2’s variance stabilizing normalization was implemented on remaining data (*25*). KEGG Orthologs were assigned with UProC (*26*) and putative taxonomic assignments at the phylum level were assigned from MMETSP taxon designations (*27*). Read counts for each KEGG orthologue were then summed over genes assigned to each taxon, resulting in phylum-level signals. These environmental sequence data are deposited in the Sequence Read Archive through the National Center for Biotechnology Information under accession no. SRP136571, BioProject no. PRJNA437978.

### Taxonomic Resolution Selection

For >5 μm transcripts, phyla were selected at the level of taxonomic resolution to compromise between clarity in overall features of the data and inclusion of the greatest number of transcriptional signals. This led to an investigation of 14 different eukaryotic phyla in the >5μm fraction. While some phyla are dominated by photosynthetic genera (e.g. the Bacillariophyta), some phyla potentially include signals from mixotrophic or heterotrophic genera (e.g. the Dinophyta), and so we conservatively use the term eukaryote throughout. In the >0.2 μm fraction, because prokaryotes have fewer genes, the compromise between taxonomic resolution and data interpretability was less severe, and so some specific taxa of known importance (*28, 29*) and interest were highlighted, such as *Prochlorococcus* (HL and LL ecotypes), *Synechococcus*, SAR11, and *Crocosphaera*, while other taxonomic groups were left at the phylum level (such as *Actinobacteria* and *Planctomycetes*). A complete list of taxa examined as well as the distribution of their diel transcriptional signals across clusters is available in Supplemental Data 4.

### Determination of Diel Periodicity

For all datasets, diel periodicity was determined using the rank-based Jonckheere-Terpstra Umbrella test as implemented in R’s RAIN package (*30*). Data were first detrended (the linear regression with respect to time was subtracted from the time series) to increase power of rhythmicity detection using the detrend function in R’s pracma package (*31*), and after RAIN implementation, the Benjamini-Hochberg FDR control procedure was implemented to assess significance at the p=0.05 level for each data type, considering data sets separately because time series between data types were not fully overlapping. For metabolite data, which was measured in triplicate, observations across all replicates were used for determination of periodicity, while the averages across replicates for each time point were used for clustering analysis. Only significantly diel signals were retained for further analysis.

### Clustering Analysis

Detrended diel time series were scaled (the time series mean was subtracted and points were divided by their standard deviations using the scale function in base R) in order to make data dimensionless and reduce the impact of magnitude on the construction of distance matrices, maximally preserving the shape of the periodic element of the time series over all other features. A distance matrix was calculated using Euclidean distance. The Hopkins statistic was calculated for this distance matrix to assess the meaningfulness of clustering and value of h=0.79 was found, indicating structure in the data which cannot be explained by a random distribution of distances between objects. To determine a well-fitting clustering method, hierarchical clustering (implemented by hclust function in the R stats package), partitioning about medoid clustering (a version of knn clustering more robust to outliers, calculated using the clara function from the R cluster package (*32*)), and training of self-organizing maps (using the R kohonen package (*33*)) were implemented. To evaluate the fit of each of these results, the Calinski-Harabasz (CH) metric (using calinhara function from R library fpc (*34*)) and average silhouette distance (using the silhouette function from R package cluster (*32*)) were calculated and compared (Supplemental Figure 1). On the basis of higher average silhouette width and CH score for all potential clusterings, SOM was selected as the clustering method for the data. We used the heuristic of identifying the ‘elbow’ in decreasing average silhouette width for initially selecting 4 as the operational number of clusters. To inspect the fits for more detail, we calculated the per-cluster average number of time series with negative silhouette widths (interpreted as misclassifications) for 3, 4, and 5 cluster clusterings. We found that 4 clusters had the fewest average misclassifications per cluster. To further compare these potential clusterings, we generated ordered dissimilarity images and silhouette profiles for the 4 and 5 cluster clusterings. Silhouette width profiles were then constructed for each cluster for more detailed inspection of cluster coherence, and 4 clusters was selected as the optimal number for these data.

**Fig. S1.**
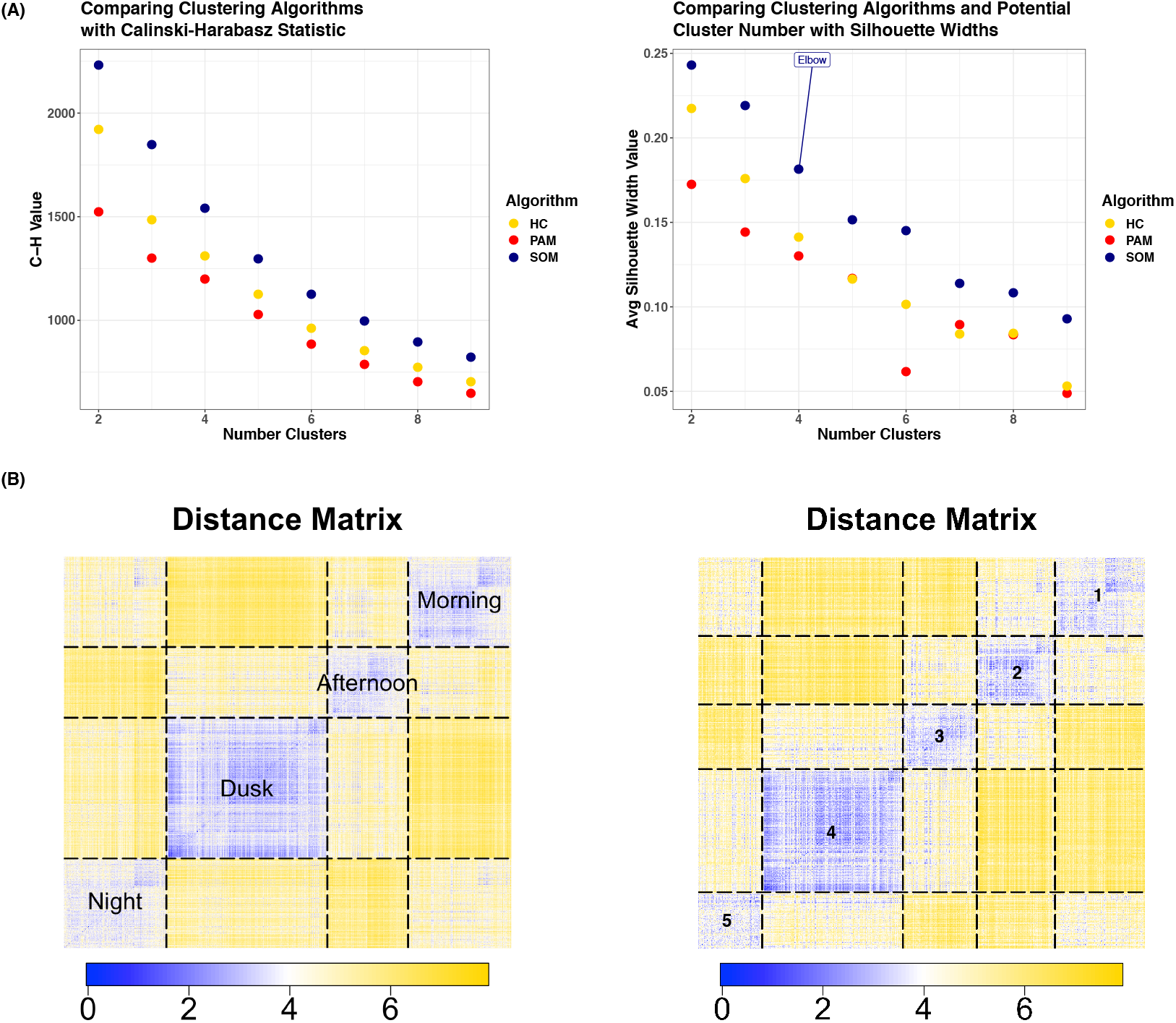
Comparison of cluster metrics for archetype clustering. Top panels show dynamics in clustering Calinski-Harabasz index and average silhouette width for increasing number of clusters comparing 3 clustering algorithms – self-organizing maps (SOM), hierarchical clustering (HC), and clustering about perimedoids (PAM). SOM was chosen for further clustering based on the advantage in C-H index and average silhouette width (A). The number of clusters was then selected based on the plateau in average silhouette width between four and five clusters. We then used ordered dissimilarity images (ODIs) to compare the four-cluster and five-cluster results (B). For additional information, silhouette profiles were constructed for all clusters in both clusterings. The 4 cluster clustering was chosen on the heuristic basis of higher maximum silhouette widths for all clusters in the 4 cluster SOM and fewer negative silhouette widths in all clusters (using negative silhouette width as a proxy for misclassification). Summary statistics for silhouette profiles are provided in Data S8. Briefly, the SOM using 4 clusters had 235/6273 (3.7%) of silhouette widths less than 0 and maximum per-cluster silhouette widths between 0.322-0.456, while the SOM using 5 clusters had 353/6273 (5.6%) of silhouette widths less than 0 and maximum per-cluster silhouette widths between 0.265-0.419, indicating fewer misclassifications in the 4 cluster SOM and greater within-cluster similarity.

### Calculation of Peak Rank Time

To estimate mean peak time for transcriptional profiles, a rank-based heuristic was calculated. For a given transcript or biomolecule, the expression levels at each measurement were ranked. The ranks from all the 0200 measurements, 0600 measurements, etc, were averaged, and the peak mean rank time was defined as the time with the highest average, where ties were summarized as the center between tied times (for example, if a transcript had the same mean rank for 0200 measurements and 0600 measurements, the mean peak rank would be defined as 0400). All peak rank time estimates along with original and rotated NMDS coordinates provided in Supplemental Data 5.

### Assessing Average Peak Rank Time Difference

For all KEGG orthologues with diel expression in at least 4 different taxonomic groups, we found all circular pairwise differences in peak times based on the peak times calculated in the above section. For example, if an orthologue had a 0600 peak time for one taxon and a 1400 peak time for another taxon, the pairwise difference would be 8 hours. The average of all of these pairwise differences was taken for each KO. To assess whether or not an orthologue had significantly small average peak-time difference (i.e., taxa tend to peak expression of this orthologue at the same time of day), we employed a monte carlo simulation using multinomial draws of size (number of taxa with diel expression of the orthologue) from the population of all observed diel transcripts 10000 times. The empirical distribution of average peak-time difference was then used as a null to calculate the simulated p-value of each orthologue. Significance was thresholded using the Benjamini-Hochberg procedure at FDR 10% (p<0.1).

### Pathway Enrichment Analysis

The 5722 transcripts identified as significantly diel were assigned to 4193 unique KEGG Orthologs (KO’s). Since many KO’s may be assigned to multiple pathways, pathways were manually curated by inspection to eliminate redundant, ambiguous, or otherwise inappropriate assignments. This resulted in 258 unique pathway assignments amongst 3097 unique KO’s; the remaining 1096 KOs could not be unambiguously unassigned to a pathway (Supplemental Data 6). Diel transcripts with an assigned KO were also mapped to taxonomic classifiers – eukaryotes, cyanobacteria (photoautotrophic bacteria), and non-eukaryotic heterotrophs (nearly exclusively bacterial, see Supplemental Data 6), and enrichment analysis was performed for each KEGG pathway for each of these groups using Fisher’s exact test. To account for multiple testing, the Benjamini-Hochberg adaptive FDR control procedure was implemented using a significance threshold of FDR=10% (p<0.1).

## Supplementary Text

### Implications of Diel Cycle on Micronutrient Dynamics at Station ALOHA

#### Cobalamin Dynamics

While the oligotrophic NPSG is nitrogen limited, micronutrients such as vitamin B12 (cobalamin) are a valuable resource to a wide array of organisms, like eukaryotic phytoplankton and heterotrophic bacteria, that need to acquire the compound exogenously (*35–37*). In eukaryotic phytoplankton and other higher organisms experiencing cobalamin limitation, S-adenosyl homocysteine (SAH) is elevated and SAH’s precursor and near-universal methyl donor S-adenosyl methionine (SAM) decreases (*38, 39*). We find SAH in the afternoon cluster while SAM peaks at dusk with POC (Supplemental File S3) This daytime increase in the SAH/SAM ratio potentially indicates a temporary bottleneck in the methionine cycle that could be due to cobalamin availability. Cobalamin-like compounds photodegrade quickly in the ocean and the turnover time has been estimated to be on the order of hours to days (*40*). We therefore looked for more evidence of diel cobalamin dynamics in our data.

Cobalamin is produced by some bacteria and archaea, and pseudocobalamin, a cobalamin analogue with a different lower ligand, is produced by cyanobacteria (*41*). These chemical variants of Vitamin B12 require enzymatic remodeling in order to complete vitamin cross feeding between these groups (*41, 42*). We found evidence of diel cyanobacterial pseudocobalamin production (Supplemental File S3). In addition, we identify *Crocosphaera* cobalt-uptake receptors (cbiMN) and cobalamin-synthesis transcripts (cobW and cobSV) in the dusk cluster, followed by a putative cobalamin transport membrane protein in the night cluster (bacA, (*43*)). *Crocosphaera* has been shown to produce and excrete a cobalamin-like compound (likely pseudocobalamin) at a high rate in culture (*44*). Notably, in this study, *Crocosphaera* transcripts involved in pseudocobalamin synthesis peak during *Crocosphaera*’s nighttime nitrogen-fixing metabolic phase. While this coupling makes intuitive sense because of the high nitrogen content of cobalamin-like compounds and likely daytime light-driven degradation, it also adds evidence supporting a mechanistic linkage between intracellular nitrogen availability and the production of cobalamin-like compounds proposed by earlier work (*45*). This connection was posited by Bonnet et al. (*44*), who found that N-replete cultures of *Synechococcus* (a cyanobacterium which does not fix nitrogen) produce more cobalamin-like compounds than N-limited cultures.

Concurrent with the nighttime *Crocosphaera bacA* transporter, we find diel cobalamin transporters (*bacA* and *btuB*) in 3 different eukaryotic phyla (Supplemental File S1). There is additional diel expression in the haptophytes specifically, with both parts of the cobalamin-dependent methionine synthesis gene (methyltetrahydrofolate-homocysteine methyltransferase *metF* and *metH*) landing in the afternoon cluster (Supplemental File S3). Though it is not significantly diel, *CobS*, which is required for lower-ligand remodeling of pseudocobalamin to cobalamin, is expressed in six eukaryotic taxa. These lines of evidence point towards active production, transport, and remodeling, and indicate that cobalamin-like compounds have diel dynamics across broad taxa in the surface ocean.

#### Iron Ligand Transporter Dynamics

Trace metals, and particularly iron, more broadly comprise another class of micronutrients which are present in the NPSG at very low levels (*46–49*). Iron is a critical metal cofactor to a wide array of enzymes (*50*), including those involved in photosynthesis and nitrogen fixation (*51*). Dissolved trace metal concentrations measured during the same cruise used for sampling in this study show no obvious diel fluctuation in trace metal concentrations and stronger day-to-day variability than dawn/dusk variability in dissolved iron concentrations (*49*). Beyond the size of the dissolved iron pool, however, microbial iron cycling is also mediated by cellular iron demand and the secretion, uptake, and exchange of organic high-affinity iron-binding ligands (e.g. siderophores). Some siderophores produced by marine heterotrophic bacteria exhibit photoreactivity and generate photoproducts that are more weakly iron-binding (*52–54*). Field observations in the NPSG have found siderophores with structural similarity to aquachelin and vibrioferrin, two siderophores with known photoreactivity (*47, 48, 54*). Photoreactivity potentially indicates diel changes in the speciation of the available iron pool for microbial uptake, and has been implicated in potential metabolic exchange between heterotrophic bacteria and eukaryotic phytoplankton (*53*). Therefore, we may also expect the demand and uptake dynamics of iron may be impacted by diel shifts in metabolic activities amongst the NPSG microbial community.

Across all analyzed taxa, metal and metal-ligand transporters make up 21% (35/168) of diel transporters. Heterotrophic bacteria (primarily alpha- and gammaproteobacteria) account for 8 of these transporters, 6 of which transport either ferric iron or iron-ligands, as well as two TonB receptors, which canonically are associated with siderophore active transport. Interestingly, all of these heterotrophic iron transporters belong to either the morning or night clusters, and the *fhuA* (ferrichrome outer membrane uptake receptor) orthologue was identified as significantly synchronous in our peak timing analysis (main text Figure 5). Ferrioxamines, a class of hydroxamate siderophores similar to ferrichrome, were also measured at the time of sampling and are shown to be the most abundant class of siderophores at the surface (*48*). Ferrioxamines have also been isolated from marine gammaproteobacteria (*48*). If we only consider the direct effects of light-forcing, the widespread diel expression of iron uptake amongst heterotrophic bacteria is unexpected because, unlike photosystem proteins of photoautotrophs, the bacteriochlorophyll *a* pigment that we find evidence for expression of requires a magnesium cofactor (*55*) while proteorhodopsin uses a retinal chromophore (*56, 57*). Therefore, we might not expect morning iron requirements to be related directly to photoheterotrophy. However, the morning cluster is significantly enriched for ribosomal subunits in heterotrophs and therefore morning may be an important proteinogenic time. Thus, diel iron uptake may be in response to demand for iron as an enzyme cofactor. For example, the afternoon cluster and morning cluster contain *Roseobacter* and SAR116 transcripts for the iron-manganese containing form of superoxide dismutase, which may be an important enzymatic accessory to photoheterotrophy (58) and protection from UV-induced ROS during cellular replication.

We can contrast heterotrophic bacteria with the cyanobacteria. For instance, in *Prochlorococcus*, we identified diel transcription of a ferric iron uptake receptor with peak timing in the afternoon, coinciding with maximum light incidence and diel transcriptional peaks of high-iron machinery such as photosystem II and ferredoxin (*59*). In *Crocosphaera*, we find ferrous iron transporters in the dusk cluster when iron demand is high for synthesizing nitrogen fixation machinery, accompanying iron storage protein bacterioferritin and FeS cluster assembly protein *sufC* (*51*). Together, we find evidence for partitioned iron demand and uptake across bacteria throughout the diel cycle – where heterotrophs express siderophore uptake receptors in the morning, a non-diazotrophic cyanobacteria expresses ferric iron uptake receptors in the afternoon, and a diazotrophic cyanobacteria expresses ferrous iron transporters (potentially indicating intracellular iron recycling) at dusk, corresponding to concurrent expression of superoxide dismutases, photosynthesis accessory proteins, and nitrogen fixation machinery, respectively.

## Supplementary Data Files

**Data S1. RAIN analysis**

Files containing the results from RAIN analysis for all analyzed transcripts, lipids, metabolites separated by data source. Diel YES/NO indicates BH adjusted p-value at 0.05.

**Data S2. Taxon analysis**

Table detailing for each taxon studied how many unique KEGG orthologs were observed, how many of them were diel, and whether they came from the small size fraction or large size fraction transcriptomes.

**Data S3. SOM Clustering**

Details for every diel signal from SOM clustering, including the signal, its cluster, the silhouette width for that signal (see methods), and the nearest neighboring cluster for that signal. Files are divided by data source.

**Data S4. Diel transcript for cluster**

Table showing the proportion of diel transcripts in each cluster for each taxon studied with diel signals.

**Data S5. NMDS ordination**

NMDS ordination results including mean peak rank time calculation. In this calculation, peak rank time works as follows: Peak rank time of 1 indicates 10pm peak, 2 indicates 2am, 3 indicates 6am, 4 indicates 10am, 5 indicates 2pm and 6 indicates 6pm. Table includes coordinates from initial NMDS ordination as well as coordinates rotated by pi/16 to align midnight peak-time with the top-center of the plot.

**Data S6. KEGG Pathway Enrichment**

Summary of KEGG pathway enrichment analysis results.

**Data S7. Rank Time Difference**

Summary of Mean Peak Rank Time Difference Analysis.

**Data S8. Clustering Summary Statistics**

Summary statistics of silhouette profiles comparing SOM clusterings with 3,4, and 5 clusters.

