## Supplemental Information for "Community-scale Synchronization and Temporal Partitioning of Gene Expression, Metabolism, and Lipid Biosynthesis in Oligotrophic Ocean Surface Waters"

**This PDF file includes:**

Materials and Methods
Supplementary Text
Figs. S1 to S2
Captions for Data S1 to S8

**Other Supplementary Materials for this manuscript include the following:**

Data S1 to S8

### Materials and Methods

**Fig. S1.**

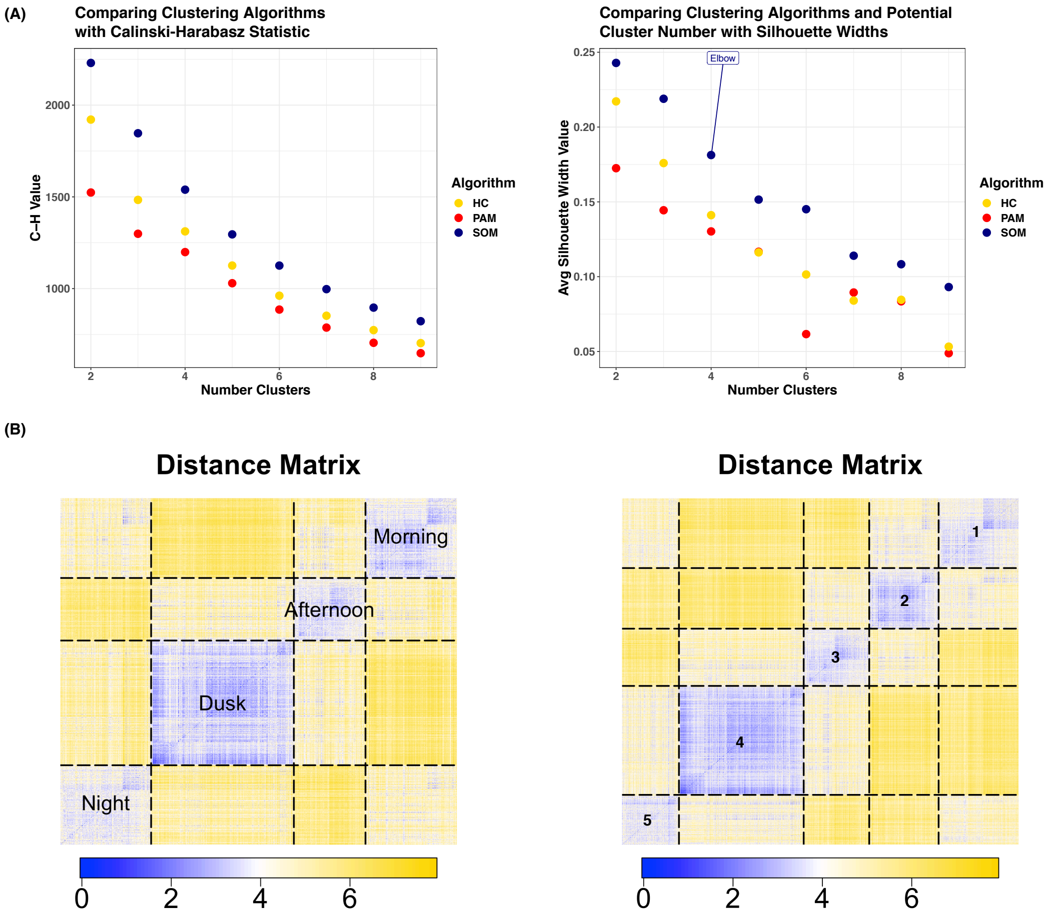


Comparison of cluster metrics for archetype clustering. Top panels show dynamics in clustering Calinski-Harabasz index and average silhouette width for increasing number of clusters comparing 3 clustering algorithms – self-organizing maps (SOM), hierarchical clustering (HC), and clustering about perimedoids (PAM). SOM was chosen for further clustering based on the advantage in C-H index and average silhouette width (A). The number of clusters was then selected based on the plateau in average silhouette width between four and five clusters. We then used ordered dissimilarity images (ODIs) to compare the four-cluster and five-cluster results (B). For additional information, silhouette profiles were constructed for all clusters in both clusterings. The 4 cluster clustering was chosen on the heuristic basis of higher maximum silhouette widths for all clusters in the 4 cluster SOM and fewer negative silhouette widths in all clusters (using negative silhouette width as a proxy for misclassification). Summary statistics for silhouette profiles are provided in Data S8. Briefly, the SOM using 4 clusters had 235/6273 (3.7%) of silhouette widths less than 0 and maximum per-cluster silhouette widths between 0.322-0.456, while the SOM using 5 clusters had 353/6273 (5.6%) of silhouette widths less than 0 and maximum per-cluster silhouette widths between 0.265-0.419, indicating fewer misclassifications in the 4 cluster SOM and greater within-cluster similarity.

**Supplementary Data Files**

**Data S1. RAIN analysis**
Files containing the results from RAIN analysis for all analyzed transcripts, lipids, metabolites separated by data source. Diel YES/NO indicates BH adjusted p-value at 0.05.

**Data S2. Taxon analysis**
Table detailing for each taxon studied how many unique KEGG orthologs were observed, how many of them were diel, and whether they came from the small size fraction or large size fraction transcriptomes.

**Data S3. SOM Clustering**
Details for every diel signal from SOM clustering, including the signal, its cluster, the silhouette width for that signal (see methods), and the nearest neighboring cluster for that signal. Files are divided by data source.

**Data S4. Diel transcript for cluster**
Table showing the proportion of diel transcripts in each cluster for each taxon studied with diel signals.

**Data S5. NMDS ordination**
NMDS ordination results including mean peak rank time calculation. In this calculation, peak rank time works as follows: Peak rank time of 1 indicates 10pm peak, 2 indicates 2am, 3 indicates 6am, 4 indicates 10am, 5 indicates 2pm and 6 indicates 6pm. Table includes coordinates from initial NMDS ordination as well as coordinates rotated by pi/16 to align midnight peak-time with the top-center of the plot.

**Data S6. KEGG Pathway Enrichment**
Summary of KEGG pathway enrichment analysis results.

**Data S7. Rank Time Difference**
Summary of Mean Peak Rank Time Difference Analysis.

**Data S8. Clustering Summary Statistics**
Summary statistics of silhouette profiles comparing SOM clusterings with 3,4, and 5 clusters.


1. S. T. Wilson *et al.*, Coordinated regulation of growth, activity and transcription in natural populations of the unicellular nitrogen-fixing cyanobacterium *Crocosphaera*. *Nat Microbiol* **2**, 17118 (2017).
2. B. P. Durham *et al.*, Sulfonate-based networks between eukaryotic phytoplankton and heterotrophic bacteria in the surface ocean. *Nat Microbiol*, (2019).
3. A. K. Boysen *et al.*, Diel oscillations of particulate metabolites reflect synchronized microbial activity in the North Pacific Subtropical Gyre. *bioRxiv*, 2020.2005.2009.086173 (2020).

4. E. Bligh, W. Dyer, A rapid method of total lipid extraction and purification. *Can. J. Biochem. Physiol* **37**, 911-917 (1959).
5. A. K. Boysen, K. R. Heal, L. T. Carlson, A. E. Ingalls, Best-matched internal standard normalization in liquid chromatography–mass spectrometry metabolomics applied to environmental samples. *Anal. Chem.* **90**, 1363-1369 (2018).
6. B. MacLean *et al.*, Skyline: an open source document editor for creating and analyzing targeted proteomics experiments. *Bioinformatics* **26**, 966-968 (2010).
7. M. Fountoulakis, H.-W. Lahm, Hydrolysis and amino acid composition analysis of proteins. *J. Chromatogr. A* **826**, 109-134 (1998).
8. K. W. Becker *et al.*, Daily changes in phytoplankton lipidomes reveal mechanisms of energy storage in the open ocean. *Nat Commun* **9**, 5179 (2018).
9. K. J. Popendorf, H. F. Fredricks, B. A. S. Van Mooy, Molecular ion-independent quantification of polar glycerolipid classes in marine plankton using triple quadrupole MS. *Lipids* **48**, 185-195 (2013).
10. J. R. Collins, B. R. Edwards, H. F. Fredricks, B. A. S. Van Mooy, LOBSTAHS: An adduct-based lipidomics strategy for discovery and identification of oxidative stress biomarkers. *Anal. Chem.* **88**, 7154-7162 (2016).
11. J. Hummel *et al.*, Ultra performance liquid chromatography and high resolution mass spectrometry for the analysis of plant lipids. *Front Plant Sci* **2**, 54 (2011).
12. C. A. Smith, E. J. Want, G. O'Maille, R. Abagyan, G. Siuzdak, XCMS: processing mass spectrometry data for metabolite profiling using nonlinear peak alignment, matching, and identification. *Anal. Chem.* **78**, 779-787 (2006).
13. C. Kuhl, R. Tautenhahn, C. Bottcher, T. R. Larson, S. Neumann, CAMERA: an integrated strategy for compound spectra extraction and annotation of liquid chromatography/mass spectrometry data sets. *Anal. Chem.* **84**, 283-289 (2012).
14. F. O. Aylward *et al.*, Diel cycling and long-term persistence of viruses in the ocean's euphotic zone. *Proc Natl Acad Sci U.S.A.* **114**, 11446-11451 (2017).
15. A. M. Bolger, M. Lohse, B. Usadel, Trimmomatic: a flexible trimmer for Illumina sequence data. *Bioinformatics* **30**, 2114-2120 (2014).
16. A. P. Masella, A. K. Bartram, J. M. Truszkowski, D. G. Brown, J. D. Neufeld, PANDAseq: paired-end assembler for illumina sequences. *BMC bioinformatics* **13**, 31 (2012).
17. N. Joshi, J. Fass. Sickle: A sliding-window, adaptive, quality-based trimming tool for FastQ files (Version 1.33) (2011).
18. E. Kopylova, L. Noé, H. Touzet, SortMeRNA: fast and accurate filtering of ribosomal RNAs in metatranscriptomic data. *Bioinformatics* **28**, 3211-3217 (2012).
19. S. M. Kiełbasa, R. Wan, K. Sato, P. Horton, M. C. Frith, Adaptive seeds tame genomic sequence comparison. *Genome res* **21**, 487-493 (2011).
20. D. R. Mende *et al.*, Environmental drivers of a microbial genomic transition zone in the ocean's interior. *Nat Microbiol* **2**, 1367-1373 (2017).
21. M. J. Harke *et al.*, Periodic and coordinated gene expression between a diazotroph and its diatom host. *ISME J* **13**, 118-131 (2018).
22. H. Li, R. Durbin, Fast and accurate long-read alignment with Burrows–Wheeler transform. *Bioinformatics* **26**, 589-595 (2010).
23. H. Alexander *et al.*, Functional group-specific traits drive phytoplankton dynamics in the oligotrophic ocean. *Proc Natl Acad Sci U.S.A.* **112**, E5972-E5979 (2015).
